# An arginine switch drives the stepwise activation of β-arrestin

**DOI:** 10.1101/2025.01.05.631353

**Authors:** Jeong Seok Ji, Yaejin Yun, Tomasz Maciej Stepniewski, Hye-Jin Yoon, Kyungjin Min, Ji Young Park, Chiwoon Chung, Ka Young Chung, Jana Selent, Hyung Ho Lee

## Abstract

β-arrestins (βarrs) play a crucial role in regulating G protein-coupled receptor (GPCR) signaling and trafficking. Canonically, interactions of βarr with the phosphorylated intracellular GPCR-tail induce a multi-step conformational transition that results in the activation of βarr. Depending on the specific interaction pattern with the receptor, βarrs adopt multiple conformational states, each tightly linked to a specific functional outcome of βarr recruitment. Despite its physiological relevance, the structural determinants of βarr activation remain poorly understood. Using a combination of molecular dynamics simulations, biochemical and cell-based experiments, we reveal how specific interactions with a prototypical GPCR promote the unbinding of the βarr2 C-tail—a crucial step in arrestin activation. Importantly, we observe that the expulsion of the C-tail is promoted by the displacement of a conserved arginine residue (Arg394) within the βarr polar core, which we dub “the arginine switch.” Our study uncovers a previously unknown molecular switch that, upon engagement, destabilizes the polar core as a crucial step in the GPCR-induced βarr activation.

## Introduction

β-arrestin1 and β-arrestin2 (referred to as βarr1 and βarr2) are multifunctional adaptor proteins that mediate the desensitization and internalization of G protein-coupled receptors (GPCRs)^1-4^. Upon agonist-induced activation of GPCRs, βarrs bind to the phosphorylated C-tails or intracellular loops of the receptor^5-10^. This leads to the recruitment of clathrin and the adaptor protein complex-2 (AP2), which triggers receptor endocytosis^11^. Subsequently, the internalized receptor can be either recycled back to the cell surface or degraded^11-13^. The interaction between βarrs and clathrin/AP2 complexes is crucial for the efficient internalization of GPCRs^1,14^. The removal of GPCRs from the cell surface through βarr-mediated internalization prevents them from transmitting extracellular signals into cells. Despite being initially identified as proteins that desensitize GPCRs, βarrs can also interact with other signaling proteins, including mitogen-activated protein kinases (MAPKs)^15,16^, Src protein kinase^17^, AKT^18^, and the NF-κB cascade^19-21^, triggering a second wave of intracellular signaling, independent of G proteins^22-24^. The development of ‘biased’ drugs, which selectively modulate the activation of GPCRs to preferentially stimulate or avoid stimulating βarrs instead of G proteins, has the potential to yield improved therapeutic outcomes with enhanced safety profiles for a broad spectrum of diseases^25^.

A hallmark of β-arrestin (βarr) is its remarkable structural flexibility. In response to interactions with GPCRs and small molecules, the protein can undergo a conformational transition in a process called activation^26^. βarr can assume multiple active conformations, which are linked to how it engages with the receptor and influence the functional outcome of βarr recruitment^27-30^. Therefore, it is crucial to understand the molecular mechanisms governing the activation of βarrs by GPCRs.

Previous structural studies have identified both inactive and active states of βarrs. The inactive state is stabilized through the autoinhibitory binding of the βarr C-terminal tail to the N-domain of βarrs^24,31-33^, which prevents the inter-domain rotation between the N- and C-domains. The autoinhibitory binding of βarr C-tail to the N-domain occludes the engagement of the receptor C-tail (hereafter referred to as the R_p_-tail). Thus, it has been hypothesized that the binding of R_p_-tail or intracellular loops of GPCRs to the N-domain of βarrs is associated with the release of βarr C-tail from the N-domain^24,34-36^. Indeed, the release of βarr C-tail from the N-domain and the subsequent inter-domain rotation have been observed in the active state of βarrs^8-10,26,37-42^.

The autoinhibitory binding of βarr C-tail to the N-domain is sustained by two distinct interfaces: the polar core and three hydrophobic elements (3E) (β-strand I and α-helix I in the N-domain and β-strand XX in the C-tail)^32,33,37,43^. The release of the βarr C-tail is a critical step in facilitating downstream signaling by exposing buried regions that are otherwise inaccessible in inactive βarr^32,44,45^. These regions include the βarr C-tail regions responsible for clathrin binding, as well as the N-domain regions of βarr that bind to scaffolding signaling kinases^14,24,46,47^. Despite the functional significance of the release of the βarr C-tail, the molecular mechanism underlying how R_p_-tail binding promotes the full displacement of the βarr C-tail remains unclear due to the absence of structural information.

In this study, we aimed to unravel initial states of βarr2 activation by a R_p_-tail. To achieve this, we simulated βarr2 in complex with a phosphopeptide (C7pp2) derived from the carboxyl terminus of CXCR7, an atypical chemokine receptor that interacts with βarrs but lacks functional coupling with heterotrimeric G proteins^48^. Our structural analysis suggests that C7pp2 can bind while the βarr2 C-tail remains attached to the N-domain of βarr2, with a partially disrupted polar core and other characteristics resembling its inactive state. These structural insights, together with biochemical and cell biological experiments, reveal the arginine switch mechanism and shed light on the activation mechanism of βarrs by GPCR C-tails.

## Results

### An intermediate state of βarr2

A previous study assessed the functional significance of CXCR7 phosphorylation in β-arrestin (βarr) trafficking and elucidated that a distal phospho-site cluster, PxPxxP (C7pp2, where P represents pSer/pThr and x is any other amino acid, Fig. 1A), significantly contributes to βarr2 recruitment and trafficking^49^. Since βarr2 is known to be stable in an auto-inhibited basal state, where its C-tail containing β-strand XX is bound to the N-domain, we hypothesized that this auto-inhibited state could only be destabilized by sufficient phosphorylation at the GPCR R_p_-tail. To verify whether the presence of 3 phosphates (pSer350, pThr352, and pSer355) in C7pp2 results in the release of βarr2 C-tail, we performed an *in vitro* clathrin binding assay using full-length βarr2^50^. The clathrin binding motif of βarr2 (residues 371-379) is only accessible for clathrin binding when βarr2 C-tail is released. Our results showed that C7pp2 binding to βarr2 does not enhance clathrin binding compared to V_2_Rpp, which effectively leads to the release of βarr2 C-tail (Fig. 1B). In contrast, in the case of C7pp3, which harbors additional phosphorylations (pSer347, pSer360, and pThr361), the βarr2 C-tail is released (Fig. 1B). These results indicate that the phosphorylation of C7pp2 at three sites is insufficient to fully displace the βarr2 C-tail.

**Fig. 1.**
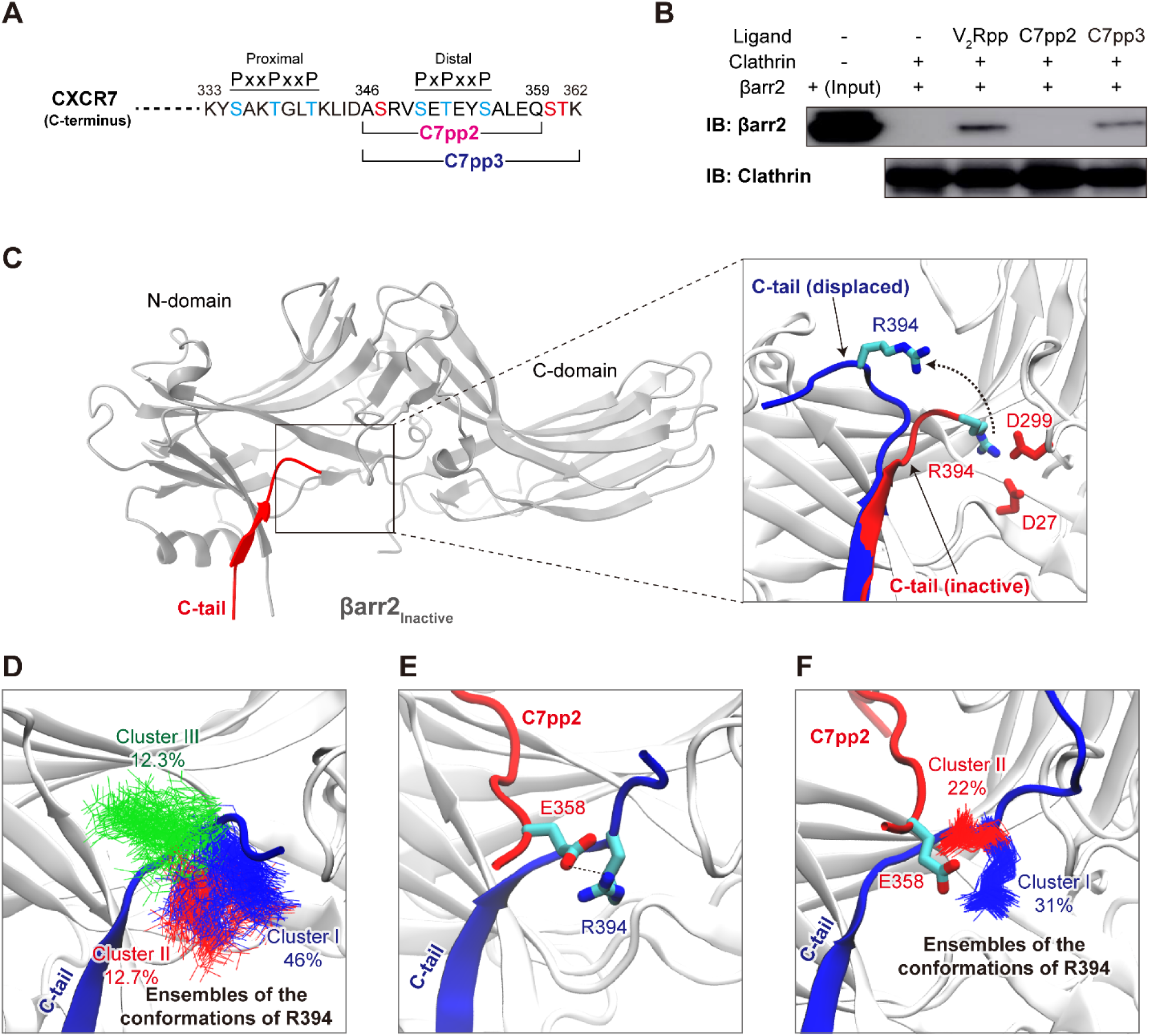
Generating a model of the intermediate state of βarr2. (A) The C-terminal sequence (residues 333-362) of CXCR7. The C7pp2 (residues 346-359) and C7pp3 (residues 346-362) peptides are indicated. Phosphorylation sites in the proximal and distal clusters (PxxPxxP and PxPxxP, where P represents pSer/pThr and x represents any other amino acid) are indicated in blue, while added phosphorylation sites on C7pp3 are denoted in red. All phosphorylated residues have been experimentally validated^57,69-74^. (B) *In vitro* clathrin binding assay for βarr2. The βarr2 binding and input are shown. Clathrin binding to βarr2 was enhanced upon binding of V_2_Rpp and C7pp3. (C) Ribbon diagram of inactive βarr2. The C-tail of the inactive βarr2 is shown in red. In an initial step of the simulation, the C-tail of βarr2 (residues 385 to 397) was manually displaced (blue) from the initial inactive conformation. (D) Representative conformations of R394 in each cluster. The frames obtained from simulations of the generated model (7 × 200 ns, restraints applied to the backbone of βarr2) were clustered based on the conformation of R394. The ensembles of the hit conformations of R394 are shown in blue (cluster I), red (cluster II), and green (cluster III) respectively. (E) The βarr2-C7pp2 complex model obtained from docking analysis. The most populated conformation of βarr2 obtained from clustering analysis was used to dock the C7pp2 peptide. (F) The representative conformations of R394 in each cluster from the simulation results. The ensembles of the hit conformations of R394 are shown in blue (cluster I) and red (cluster II). The βarr2-C7pp2 complex model obtained from docking result was used for simulations (7 × 300 ns, last 50 ns taken for analysis).

To better understand the intermediate state where the βarr2 C-tail is not fully displaced when bound to C7pp2, we used a sequential modeling protocol of molecular dynamics simulations. First, we simulated the re-binding of the C-tail, starting from an unbound conformation, to explore intermediate conformational states of the C-tail (7 × 200 ns, Fig. 1C). Our analysis specifically focused on Arg394 within the distal C-tail, as previous experimental findings have underscored the conformational rearrangement of this residue as an important step in C-tail displacement^50^. In the inactive conformation of βarr2, Arg394 is stabilized by interactions with Asp27 and Asp299 (Fig. 1C). Monitoring the distance between Arg394 and Asp299 revealed that, in the majority of the simulation replicates, we observed spontaneous inactivation of the partially displaced distal C-tail (Fig. S1, Movie S1). Interestingly, when clustering the conformational landscape explored by Arg394 during these transition dynamics (Fig. 1D), we observed a highly populated cluster (cluster I, 46%) of conformations where Arg394 is extended away from the βarr2 surface. The relatively high population of this conformation suggests that it represents an important intermediate state explored by the distal C-tail during binding/unbinding. To verify whether such a conformation can accommodate interactions with an R_p_-tail, we docked the phosphorylated C7pp2 to the representative βarr structure of cluster I. Intriguingly, the docking algorithm generated a pose in which Glu358 of C7pp2 forms interactions with the partially displaced Arg394 of βarr2 (Fig. 1E). Thus, our structural model suggests that Glu358 of C7pp2 stabilizes the displaced Arg394 of βarr2 and, in doing so, supports the initial stage of distal C-tail displacement in an intermediate active state of βarr2 (βarr2_IM_).

To further study the obtained βarr2_IM_-C7pp2 complex, we modeled the entire distal C-tail, which is typically not resolved in experimental structures due to its high flexibility, and carried out MD simulations for structural relaxation (Fig. 1F). Intriguingly, clustering analysis of the resulting simulation runs revealed that Arg394 of βarr2 primarily adopts conformations where it maintains electrostatic interactions with Glu358 of C7pp2 (Fig. 1F).

### Two distinct binding motifs between βarr2_IM_ and C7pp2

A more detailed analysis of the interface between C7pp2 and βarr2_IM_ reveals two attachment points where C7pp2 forms polar interactions with positive residues on the βarr2_IM_ surface, referred to as site I and site II (Fig. 2A). To gain insights into the conformational flexibility of the generated complex, we conducted unrestrained MD simulations with five replicates, each running for 300 ns (Fig. S2). When analyzing individual snapshots over the simulation time, we found that the distal C-tail (blue), despite being highly flexible, stacks on top of C7pp2, thereby maintaining frequent interactions with it (Fig. S3). Additionally, we observed differences in the stability of the main contact sites I and II of C7pp2. Residues within site II explored a wider range of conformations compared to site I (Fig. S3). This variability is also reflected in the frequency of contacts established between C7pp2 and βarr2_IM_. For site I, we found very stable electrostatic interactions between the phosphorylated Ser350 in C7pp2 and Arg77 (81%), Arg148 (86%), and Arg166 (80%) in βarr2_IM_ (Fig. 2A). Furthermore, the phosphorylated Thr352 in C7pp2 contributes to the stability of site I by interacting with Arg166 (38%) of βarr2_IM_. Interestingly, for site II, our simulations revealed transient interactions between Glu358 of C7pp2 and Arg394 of βarr2_IM_, which can break and reform multiple times during a single simulation run, achieving a total stability of 32% within the simulation frames (Fig. 2A).

**Fig. 2.**
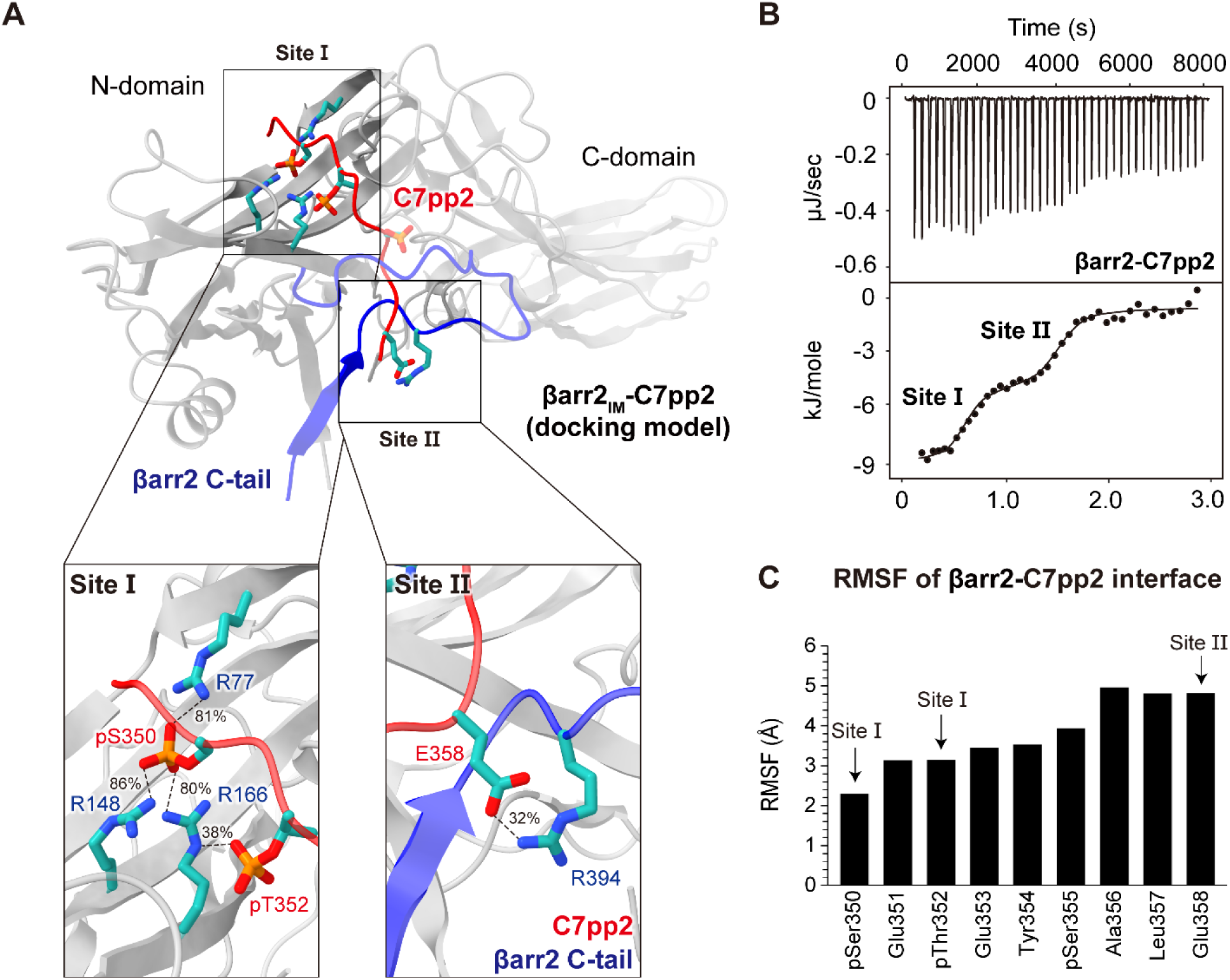
Model of the intermediate βarr2-C7pp2 state. (A) Structural model of the intermediate βarr2-C7pp2 complex. Interactions between C7pp2 and clusters of positive residues in βarr2 have been categorized into site I and site II, respectively. (B) ITC analysis of the binding of C7pp2 to βarr2. Purified βarr2 proteins were incubated with increasing concentrations of C7pp2 peptide, and the binding parameters were calculated based on the dose-response curve. βarr2 induces biphasic binding by C7pp2. (C) RMSF analysis of the Cα atoms of the simulated C7pp2 peptide. Residues of C7pp2 belonging to sites I or II are indicated with arrows.

Notably, the structural model is supported by isothermal titration calorimetry (ITC) experiments (Fig. 2B). In ITC, βarr2 exhibits a biphasic binding pattern with C7pp2, and data fitting to a two-site binding model indicates that C7pp2 contains two distinct binding motifs for βarr2. These data suggest that a primary binding site is initially occupied at lower concentrations, while a second binding site is occupied at higher concentrations during ITC titration. In this context, our structural model (Fig. 2A) suggests that the primary ITC binding site (with higher binding affinity) corresponds to site I in our βarr2_IM_-C7pp2 complex, which features abundant electrostatic interactions, whereas the second ITC binding site (with significantly lower affinity) corresponds to site II. This is further supported by the RMSF profile obtained from MD simulations (Fig. 2C), which reveals greater binding stability for site I (lower RMSF values) compared to site II (higher RMSF values).

To provide experimental evidence supporting our structural model, we disrupted the N-terminal and C-terminal interactions of C7pp2 with βarr2_IM_ using R166A (site I) and R394A (site II) mutants. Remarkably, these mutations abolished the binding peaks characteristic of site I and site II, respectively (Fig. S4). Our results indicate that C7pp2 binds to βarr2 through two distinct binding events: first, the high-affinity binding of the C7pp2 N-terminus at site I (with a *K*_D_ of 1.1 μM for the R394A mutant), and second, the low-affinity binding of the C7pp2 C-terminus at site II (with a *K*_D_ of 16.4 μM for the R166A mutant).

To further validate the contributions of pSer350 and pThr352 in the binding of C7pp2 to βarr2_IM_ at site I of our structural model (Fig. 2A), we substituted these residues with non-phosphorylated Ser350 and Thr352 and assessed the effects of each mutation using ITC (Fig. S4). Importantly, we found that the site I binding peak was abolished, confirming the crucial role of pSer350 and pThr352 in site I binding. Taken together, we conclude that C7pp2 binds to βarr2 through two distinct binding interfaces, while the βarr2 C-tail remains attached to the N-domain of βarr2.

### Partially disrupted polar core in βarr2_IM_ by C7pp2

Notably, our structural model suggests the existence of an intermediate state with interactions between Glu358 of C7pp2 and Arg394 of the βarr2 polar core (Fig. 3A). Strikingly, this structural arrangement seems to stabilize a partially displaced C-tail conformation. It is widely recognized that the polar core interactions and 3E, namely β-strand I and α-helix I in the N-domain and β-strand XX in the C-tail, play crucial roles in stabilizing the basal state of βarr2^31,33^. Consequently, both polar core and 3E must be disrupted in an active state of βarr2, which is the current conceptual framework of GPCR-βarr2 interaction. In this respect, our structural model suggest that the polar core can be partially destabilized, while the 3E interaction remains intact (Fig. S5).

**Fig. 3.**
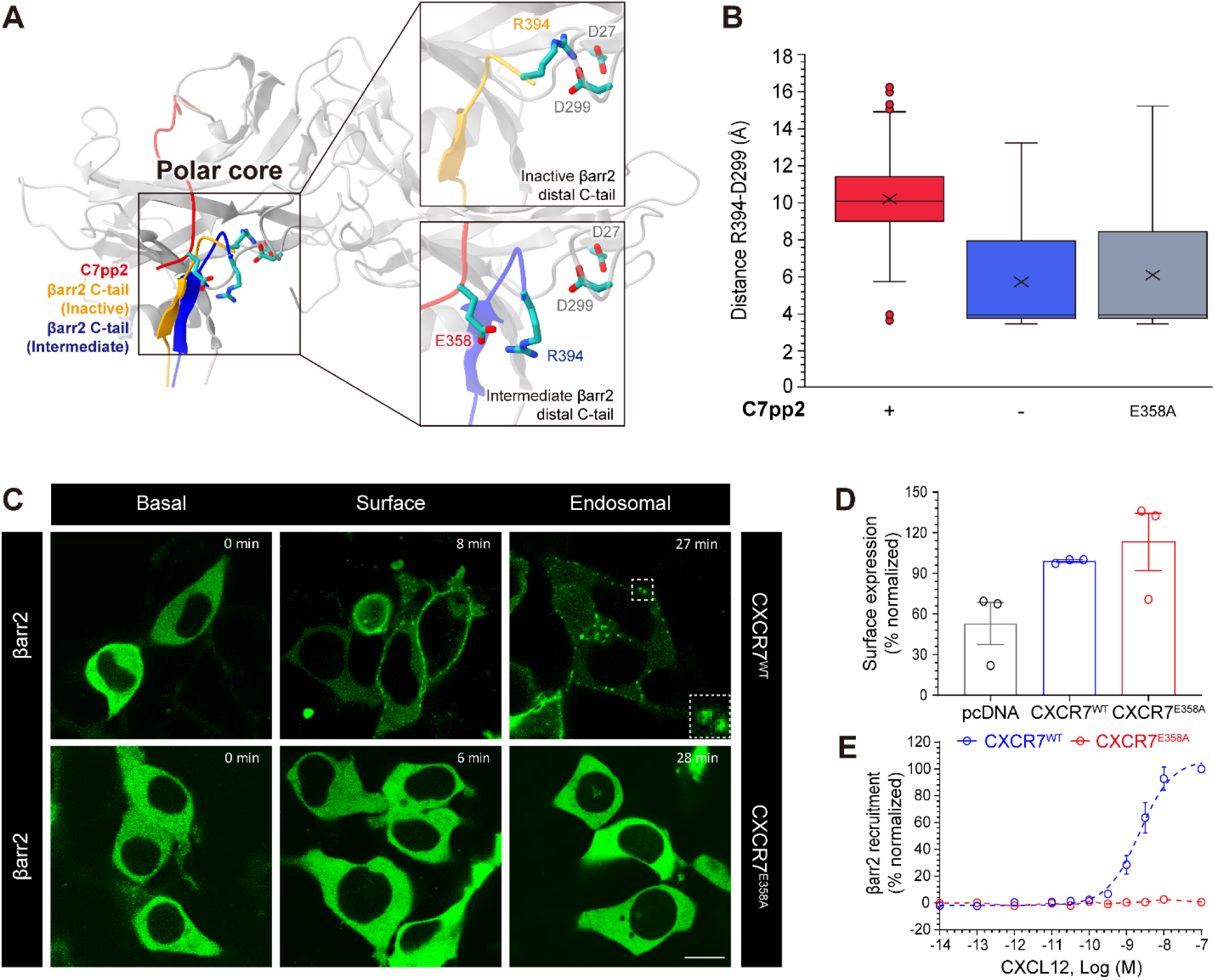
Destabilization of the polar core by C7pp2. (A) βarr2 polar core observed in the intermediate (dark blue) and inactive (orange) states of βarr2. (B) The distance between D299 and R394 (forming polar interactions in the inactive βarr2) monitored within MD simulations (5 × 300 ns). (C) CXCL12-induced trafficking of βarr2 as monitored using confocal microscopy in HEK293 cells expressing CXCR7^WT^ and CXCR7^E358A^. Scale bar is 10 μm. (D) The whole-cell ELISA-based assay to measure the surface expression of CXCR7. (E) CXCL12-induced βarr2 recruitment to CXCR7^WT^ and CXCR7^E358A^ in the Tango assay. CXCR7^E358A^ is significantly compromised in inducing βarr2 trafficking upon CXCL12 stimulation.

The polar core of βarr2 in the basal state (Fig. S5B, left panel) is comprised of charged residues from N-domain (Asp27 and Arg170), C-domain (Asp292 and Asp299), and C-terminal tail (Arg394), bringing different parts of βarr2 together, i.e., the N-domain, C-domain, and C-terminal tail hold together to prevent inter-domain rotation, which is a landmark structural change during βarr2 activation^43^. Among them, Arg394 points to the deep cleft between N- and C-domain, forming a central salt bridge, restrained by Asp27 (N-domain) and Asp299 (gate loop). It is interesting to note that Arg394 is the only residue that directly bridges all three parts (N-domain, C-domain, and C-terminal tail) among the five residues of the polar core.

To computationally verify the stabilizing effect of C7pp2 on the intermediate state, we monitored the integrity of the polar core by measuring the distance of Arg394 to Asp299 of βarr2_IM_ (Fig. 3B). While small values indicate an intact polar core (i.e., inactive βarr2) large distances reflect polar core disruption (i.e., intermediate βarr2). Interestingly, we find in the βarr2_IM_-C7pp2 complex that the polar core remains largely disrupted as indicated by an average large distance of 10 Å (Fig. 3B). In contrast, when removing the C7pp2, we can observe the re-formation of a salt bridge between Arg394 to Asp299 of βarr2 in several replicates (Fig. 3B, average distance 6 Å). This suggests that the binding of C7pp2, specifically the interaction of Glu358 in C7pp2 with Arg394 is a critical element to stabilize a broken polar core and in turn an intermediate βarr2 state.

To further corroborate this finding, we performed an *in silico* mutation of Glu358 to alanine (E358A) in the C7pp2. Remarkably, we found that this mutant also shows βarr2 inactivation in several replicates starting from a partially displaced C-tail (Fig. 3B, average distance 6 Å) emphasizing the role of Glu358 of C7pp2 and Arg394 of βarr2 interactions in stabilizing the intermediate state. To experimentally verify this observation, we conducted ITC experiments, which showed that binding at site II was indeed lost in the C7pp2^E358A^ mutant as well as in the βarr2^R394A^ mutant (Fig. S4, lower panel). Moreover, by using a confocal microscopy and Tango assay, we observed that the CXCR7^E358A^ mutant exhibited a substantial reduction in βarr2 recruitment and trafficking (Fig. 3C-E), supporting its functional role. Altogether, these results strongly suggest that the destabilization of the polar core is initiated by the formation of a salt bridge between Glu358 of C7pp2 and Arg394 of βarr2_IM_.

### HDX profile changes of βarr2 upon C7pp2 binding

To further study the conformational dynamics of βarr2 upon C7pp2 binding in solution, we adopted hydrogen/deuterium exchange mass spectrometry (HDX-MS) (Fig. 4). HDX-MS monitors the exchange between hydrogen atoms in the peptide bond backbone and deuterium in the solvent^51^. The exchange rate is affected by the conformational dynamics (i.e., exposure to the buffer and conformational flexibility) of the local area of the proteins^51^. Upon co-incubation with C7pp2, βarr2 showed increased HDX at the gate loop (Fig. 4, peptides 292-302), which is likely due to the disruption of the interaction between the gate loop (Asp299) and C-tail (Arg394). Unfortunately, we could not obtain HDX-MS data from the distal C-tail region so that we could not investigate its conformational changes. However, the HDX profile of N-terminal part of βXX was increased (Fig. 4, peptides 382-389) potentially due to increased conformational dynamics allosterically transmitted from the breakage between Asp299 and Arg394. This result supports our hypothesis that the disruption of the interaction between the gate loop (Asp299) and C-tail (Arg394) by C7pp2 further destabilizes β-strand XX. The HDX rates were decreased in the βVI through middle loop, lariat loop, and back loop (Fig. 4, 119-133, 283-291, and 305-317). Overall, our results suggest that C7pp2 binding affects primarily parts of the βarr2 C-tail through its partial displacement as well as the dynamics of the gate loop through polar core disruption.

**Fig. 4.**
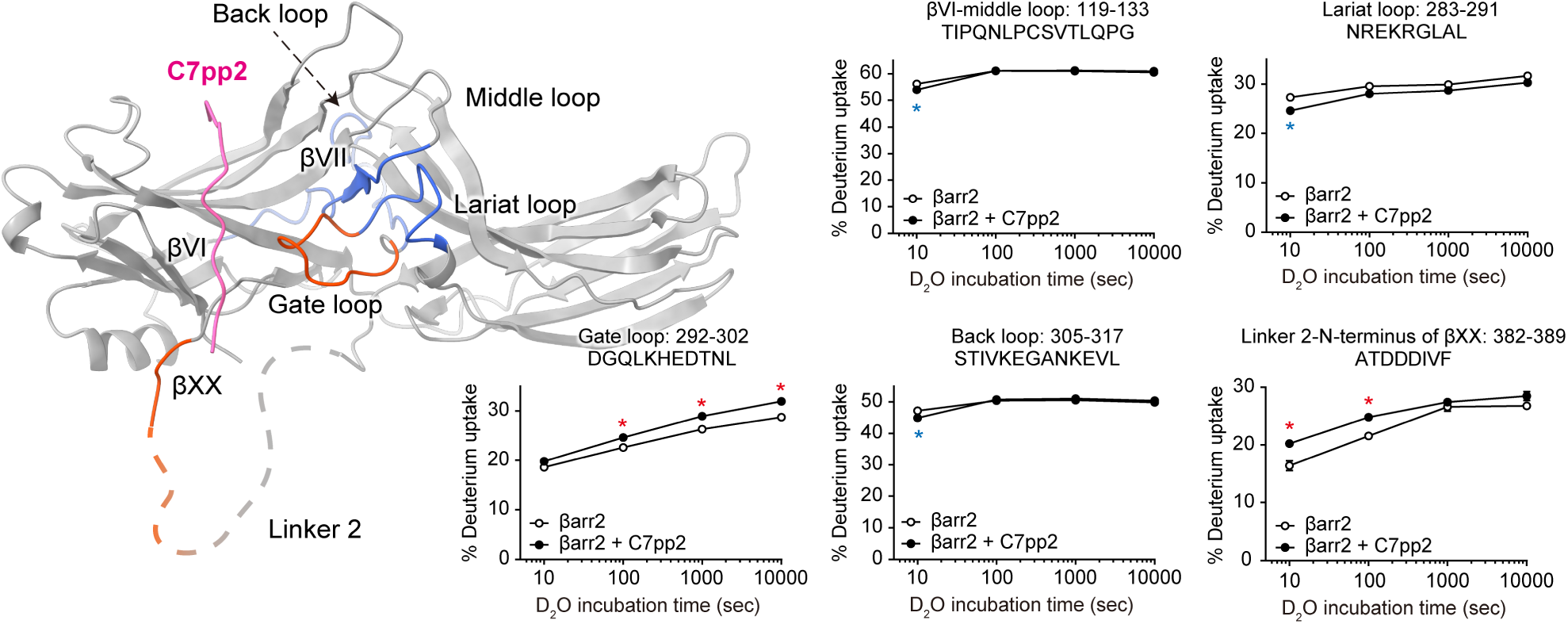
HDX profile changes of βarr2 upon co-incubation with C7pp2. Regions with increased or decreased HDX upon co-incubation with C7pp2 are colored red or blue, respectively, on the structure of the βarr2_IM_-C7pp2 complex and the HDX profiles of the corresponding peptides are shown as graphs. Data represent the mean ± standard error of three independent experiments. Statistical analysis was performed using Student’s t-test (∗p < 0.05 compared with βarr2 alone). Differences smaller than 0.2 Da were not considered significant.

### Discussion and conclusion

To activate βarr by GPCRs, interactions between GPCRs and βarr occur through two distinct interfaces. The first interface involves the binding of the receptor’s R_p_-tail to the N-domain of βarr. The second interface involves the interaction between the receptor’s transmembrane helices and cytoplasmic loops, also known as the receptor core, with the central crest loops of βarr^24,34,35,52^. In this regard, there are at least three distinct binding modes between GPCRs and βarrs, which can be mediated by both the receptor core and R_p_-tail, R_p_-tail only, or the receptor core only. It has been demonstrated that the receptor core and R_p_-tail can independently stimulate βarr activation, and these interactions trigger the release of the sequestered βarr C-tail and inter-domain rotation between the N- and C-domains^52^. One of the missing pieces of the βarr activation mechanism is how the sequestered βarr C-tail is released via interaction with GPCRs, which is considered to be the rate-limiting step^53,54^.

The previously determined structures of βarr in complex with R_p_-tails were limited to the inactive and active states of βarr. Consequently, it was not possible to directly observe how the polar core can be destabilized, as it was either in a stable or disrupted state in those structures. In this study, we aimed to investigate the process of initial βarr activation upon binding to the receptor R_p_-tail with MD simulations. The model of the intermediate state revealed several novel features that differentiate it from existing βarr structures bound to various R_p_-tails, providing valuable structural insights into the activation mechanism of βarr by the receptor R_p_-tail.

Firstly, we observed a direct binding between Glu358 of C7pp2 and Arg394 of the βarr2 C-tail. This interaction pulls the βarr C-tail away from the polar core resulting in partial disruption of the polar core through the Arg394 displacement.

Secondly, we found that C7pp2 binds to βarr2 while its C-tail remains bound to the N-domain in the proposed intermediate βarr2-C7pp2 complex. This differs from the canonical binding mode of R_p_-tails, which involves the displacement of the βarr2 C-tail. As a result, we observed two distinct binding motifs between βarr2 and C7pp2, namely the canonical binding site I and the newly discovered site II, which includes the Arg394 interaction (Fig. 2A). This observation is significant as it captures an intermediate activation state that cannot be observed in the existing active βarr2 structures. Consistently, the ITC results support the conclusion that the binding affinity of site I is significantly higher than that of site II, indicating that the binding of C7pp2 will occur sequentially from site I to site II (Fig. 2B).

Thirdly, despite the partial disruption of the polar core, the 3E interaction is still maintained. This may be attributed to the absence of two additional phosphorylations at Ser360 and Thr361, which follow the distal phospho-site cluster PxPxxP of C7pp2 (Fig. 1A). If phosphorylated Ser360 and Thr361 residues were present at the C-terminus of C7pp2, they would reside in close proximity to the 3E interaction site and disrupt it, as observed in many other active state βarr structures^37,55,56^. Indeed, V_2_Rpp, which possesses sufficient phosphorylation sites at its C-terminus, efficiently breaks the 3E interaction and triggers the release of βarr C-tail (Fig. 1B). Consistently, the clathrin binding assay confirmed that C7pp3, which contains phosphorylations on Ser360 and Thr361 similar to V_2_Rpp (Fig. 1A), enables the release of βarr C-tail (Fig. 1B). The phosphorylations on Ser360 and Thr361 appear to be mediated by GRK2 and may have an independent role, as they differ from the proximal site phosphorylated by GRK5 or the distal site phosphorylated by both GRK2 and GRK5^57^.

Based on these results, we propose the arginine switch model, in which Arg394 binds to C7pp2 to induce βarr activation (Fig. 5). In the inactive state, the arginine switch is part of the polar core, maintaining the interaction, and the βarr C-tail interacts with the N-domain through the 3E interaction. When CXCR7 R_p_-tail binds to βarr through sites I and II, the arginine switch rotates and directly binds to Glu358 of CXCR7 R_p_-tail, partially disrupting the polar core. This step represents an intermediate state before βarr becomes fully active, and the 3E interaction is still maintained. The phosphorylations of Ser360/Thr361 contribute to breaking the 3E interaction, leading to the release of the βarr C-tail, resulting in βarr folding into a fully active state.

**Fig. 5.**
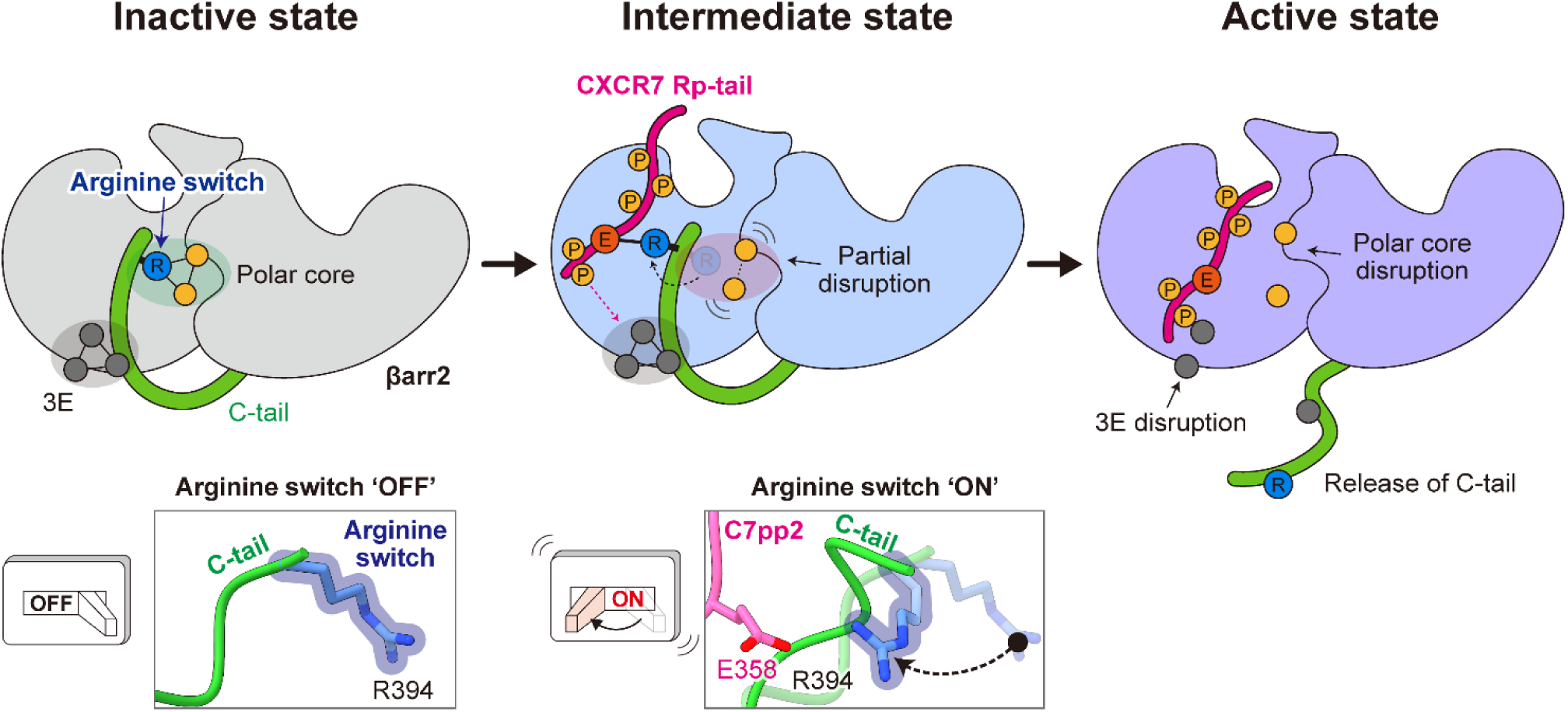
Proposed arginine switch model for βarr2 activation by the CXCR7 R_p_-tail. In the inactive state, Arg394, which we name as an “arginine switch,” is an integral component of the polar core, facilitating interaction. Simultaneously, the βarr2 C-tail engages with the N-domain through a 3E interaction. Upon C7pp2 binding to βarr2, the arginine switch undergoes rotation and forms a direct interaction with Glu358 of C7pp2, leading to partial disruption of the polar core. This stage represents an intermediate state preceding βarr2 activation, where the 3E interaction remains intact. Phosphorylation of Ser360/Thr361 further contributes to the disruption of the 3E interaction, triggering the release of the βarr2 C-tail, ultimately leading to the activation of βarr2.

While our experiments elucidate the activation mechanism of βarr2 by the receptor R_p_-tail, further investigations will be necessary to fully understand the activation mechanism of βarr. Firstly, although we explored the activation mechanism of βarr by the receptor R_p_-tail, our study did not investigate the contributions of the core region of GPCRs to βarr activation^7,52,58,59^, thus requiring further structural studies of βarr in complex with full-length GPCRs. We hypothesize that the GPCR core might also contribute to accelerating the destabilization of the polar core and 3E interactions. Alternatively, the binding of the GPCR core to βarr might trigger additional conformational changes in βarr. Secondly, the possibility of βarr activation by phosphorylated intracellular loops of GPCRs should be considered. In the case of D2-like dopamine receptors, which have shorter C-tails compared to other GPCRs, it has been shown that phosphorylated intracellular loops can interact with βarr and trigger its activation^60,61^. Thirdly, it is necessary to investigate the biological roles of the previously characterized proximal phospho-site cluster (C7pp)^40^ and the distal phospho-site cluster (C7pp2). In the case of PTH1R in living cells, βarr2 has been found to engage with both the distal phospho-site cluster and proximal phospho-site cluster, resulting in two distinct complex forms known as the tail-engaged ‘hanging’ complex and the ‘core’ complex, respectively^62^. Similarly, it is conceivable that βarr2 could adopt distinct conformations depending on the phospho-site clusters of CXCR7 while associating with the same GPCR, leading to a diverse range of functional outcomes that necessitate further investigation. Fourthly, it is possible that the intermediate state of βarr may have its own functional roles in cells, extending beyond solely serving as an activation intermediate state. In a recent report, we demonstrated that the inactive state of βarr can act as a scaffold protein and mediate its functional role^63^. Therefore, uncovering the potential biological significance of this complex, if any, awaits further investigation in future studies.

As our understanding of G protein-biased or βarr-biased ligand mechanisms expands, we become increasingly aware of their potential therapeutic benefits^64^. While βarr-biased signaling produces positive effects, G protein-dependent signaling can lead to side effects, as observed with carvedilol for both β_1_AR and β_2_AR subtypes^65,66^, [Sar^1^, D-Ala^8^] angiotensin II (TRV120027) for angiotensin II type 1 receptor^67^, and PTH for PTH1 receptor^15^. In this case, βarr-biased ligands offer a promising avenue for the discovery of novel drugs, potentially associated with reduced side effects. Our elucidation of the βarr2-C7pp2 complex provides structural insights into the intermediate state through which βarr is activated, thus laying the foundation for pharmacological studies on βarr-biased signaling.

## Data availability

The MD simulation data will be publicly available as of the date of publication at the GPCRmd repository (www.gpcrmd.org) (https://www.gpcrmd.org/dynadb/publications/1525/).^68^

## Author contributions

J.J., Y.Y., and H.H.L. conceived and designed the experiments. J.J. and Y.Y. performed the biochemical experiments. T.S. and J.S. designed and performed computational experiments including molecular modelling, docking, and molecular dynamics simulations. J.Y.P., C.C., and K.Y.C. performed the HDX-MS experiments. J.J., Y.Y., and H.H.L. analyzed the data and wrote the manuscript, with contributions from all authors. All the authors reviewed and edited the manuscript.

## Ackonwledgements

This study was supported by grants from the National Research Foundation (NRF) of Korea funded by the Korean government (2022R1A2B5B02002529, 2022R1A5A6000760, and 2021R1A2C3003518) and by the Bio & Medical Technology Development Program of the National Research Foundation (NRF) funded by the Korean government (MSIT) (RS-2024-00396026). T.M.S was supported by resources from Sara Borrell grant CD22/00007 funded by the Institute of Health Carlos III (ISCIII), resources of grant 2021 SGR 00046 funded by Agència de Gestió d’Ajuts Universitaris i de Recerca Generalitat de Catalunya (AGAUR) and the National Center of Science, Poland, grant 2017/27/N/NZ2/02571. J.S. acknowledges funding from MICIU/AEI/10.13039/501100011033 and the ERDF/EU (grant number PID2022-137161OB-I00). J.S. acknowledges further funding from the Horizon Europe Project OBELISK under the grant agreement 101080465. We thank Dr. Mithu Baidya and Parishmita Sarma from IIT Kanpur for the Tango assay.

