## Supplementary information for "An arginine switch drives the stepwise activation of β-arrestin"

### Materials and methods

#### Molecular dynamics simulation and data analysis

For MD simulation, we have started from the structure of βarr2 in an inactive conformation (PDB: 3P2D)[1]. The structure was curated including the modelling of missing residues and protonation state assignment using the MOE package (www.chemcomp.com). To match experimental conditions, the sequence of the structure was reverted to that of a rat (Uniprot code: P29067).

To simulate rebinding of the distal C-tail (Fig. 1D), we have modelled the βarr2 C-tail starting from residue 395 to residue 397 and then manually displaced the C-terminal part of the tail from the core of βarr2. Afterwards the system was simulated in 7 replicates for 200 ns, allowing the C-tail to spontaneously assume an inactive conformation (positional constraints applied to backbone atoms of βarr2 residues until residue 388, allowing C-tail flexibility). The frames were clustered using the internal algorithm in VMD[2] based on the RMSD of the Arg394 heavy atoms (cutoff 3.7).

The intermediate βarr2-C7pp2 complex was generated by docking the peptide into a representative conformation extracted from the main cluster (Fig. 1D). The peptide (residues 348-359) was docked using the HADDOCK 2 web server[3]. The binding site was defined using positions within βarr2 canonically associated with GPCR C-tail binding. The final pose was selected based on scoring as well as guidelines from available βarr-C-tail complexes. Subsequently, the whole βarr2 C-tail was modelled, and the system was allowed to relax (7 × 300 ns, positional constraints applied to backbone atoms of βarr2 residues until residue 388 as well as the docked peptide backbone). Aiming to analyze only frames where the system was equilibrated, the last 50 ns of each run was taken into account. The resulting frames were clustered based on the RMSD of the Arg394 heavy atoms (cutoff 1). From these frames, we extracted a βarr2-C7pp2 conformation for subsequent simulation runs. When studying the stability of the βarr2-C7pp2 complex, as well as the impact of the E358A mutant and lack of the C7pp2 tail (Fig. 3B) the systems were simulated in 5 replicates for 300 ns.

All generated systems were solvated (using TIP3P model water) and neutralized with NaCl ions (0.15 M concentration) using the CHARMM-GUI web server[4]. System parameters were obtained from the Charmm36M forcefield[5,6]. Simulations were carried out using Acemd3[7]. Each system initially underwent 20 ns of equilibration in conditions of constant pressure (NPT) with positional constraints applied to the backbone atoms. For production runs the systems were simulated in conditions of constant volume (NVT) using a 4 fs timestep. Simulations were analyzed using VMD[2]. Simulations are made available at the online GPCRmd resource[8]: (https://www.gpcrmd.org/dynadb/publications/1525/)

**Cloning, protein expression, and purification**

CXCR7 phosphopeptides (C7pp2 and C7pp3) for clathrin binding assay and ITC experiments were synthesized from NovoPep and Tufts University Core Facility (Fig. 1A). The *Rattus norvegicus* wild-type βarr2 was cloned into the pET-28a expression vector and then were transformed into *Escherichia coli* BL21(DE3)pLysS cells (Invitrogen). Cells harboring the plasmids were grown in Luria Bertani (LB) broth containing 70 μg $mL^{-1}$ chloramphenicol and 50 μg $mL^{-1}$ kanamycin until the optical density (at 600 nm) reached 1.0–1.2 at 37°C. Further, protein expression in the cells was induced by using 0.1 mM isopropyl-β-D-1-thiogalactopyranoside (IPTG), after which the cells were incubated for 16 h at 16°C.

To isolate the βarr2 protein fused to an N-terminal $His_6$-tag, cells were collected by centrifugation at 4,000 × g at 4°C for 10 min and resuspended in buffer A (20 mM Tris-HCl pH 8.0, 500 mM NaCl, and 5 mM imidazole) supplemented with 1 mM phenylmethanesulfonylfluoride (PMSF). The cells were lysed by passing through a

Microfluidizer (Microfluidics, Westwood, MA, USA), and the cell lysate was cleared by centrifugation. The supernatant was collected and loaded onto a Ni-sepharose affinity column (GE Healthcare, Little Chalfont, UK) and washed with buffer A. The protein was eluted using buffer A supplemented with a gradient of imidazole concentrations ranging from 100 mM to 1 M. The eluted fractions were desalted into buffer B (20 mM Tris-HCl pH 8.0 and 5 mM β-mercaptoethanol) containing 100 mM NaCl. Subsequently, further purification was carried out using affinity chromatography with a HiTrap heparin column (GE Healthcare). The proteins were eluted from the heparin column using buffer B supplemented with 1 M NaCl. To achieve further purification, eluted βarr2 was concentrated and subjected to gel filtration using a HiLoad 16/60 Superdex 200 prep-grade column (GE Healthcare). The column was pre-equilibrated with buffer B containing 200 mM NaCl and protein-containing fractions were collected. Protein was concentrated to approximately 12 mg $mL^{-1}$. The protein concentration was determined by measuring the absorbance at 280 nm.

**Clathrin binding assay**

To evaluate the binding of clathrin to wild-type βarr2, a clathrin binding assay was conducted in the absence or presence of a 5:1 (peptide:βarr2) molar ratio of ligand ($V_2$Rpp, C7pp2, and C7pp3). For each condition, 20 μg of βarr2 was incubated with the respective peptide in a binding buffer consisting of 20 mM Tris-HCl pH 8.0, 100 mM NaCl, and 1 mM DTT, at 4°C for 1 h. Following the incubation, the volume of each mixture was adjusted to 100 μL using the binding buffer. Subsequently, 30 μL of GST beads, containing 30 μg of GST-clathrin, were added to the mixture, which was then agitated for 2 h at 4°C. The GST beads with bound GST-clathrin were centrifuged at 20,000 × g. The beads were subsequently washed with 1 mL of the binding buffer 5 times. Following the washing steps, the beads were incubated with 50 μL of 25 mM GSH buffer. Clathrin binding to βarr2 was measured by Western blot analysis with an anti-his-tag mouse mAb (1:5000, MBL). Secondary antibodies used were anti-mouse IgG, HRP-linked Antibody (Cell Signaling Technology). Blots were re-probed with a GST mouse mAb (1:5000, Cell Signaling Technology) to ensure equal loading of GST-clathrin for each reaction.

**Isothermal titration calorimetry (ITC)**

ITC experiments were performed using Affinity ITC instruments (TA Instruments, New Castle, DE, USA) at 298 K. 200 μM of wild-type βarr2, which was prepared in a buffer containing 20 mM HEPES pH 7.0 and 200 mM NaCl was degassed at 298 K before measurements. Using a micro-syringe, 2.5 μL of 1 mM C7pp2 peptide solutions was added at intervals of 200 s to the wild-type βarr2 solution, in the cell with gentle stirring. The ITC experiments for the βarr2 mutants (R166A and R394A) and the C7pp2 mutants (C7pp2-1, C7pp2-2, and $C7pp2^{E358A}$) (Fig. S4) were conducted under the same running conditions.

**Surface expression assay**

To study the receptor surface expression of $CXCR7^{WT}$ and $CXCR7^{E358A}$, a whole cell-based receptor surface ELISA was performed as previously discussed[9]. Briefly, cells transiently transfected with either $CXCR7^{WT}$ or $CXCR7^{E358A}$ were seeded in a 24-well plate (corning) pre-coated with 0.01% poly-D-Lysine at a density of 0.2 million cells per well and incubated at 37°C for 24 h. After 24 h, the plate was taken out and the growth media was removed. The cells were once washed with ice-cold TBS followed by fixation with 4% Paraformaldehyde (w/v in TBS) on ice for 20 min. Thereafter, cells were washed 3 times (400 μL in each wash) with TBS followed by blocking with 1% BSA prepared in TBS at room temperature for 90 min. Subsequently, the cells were incubated with anti-FLAG M2-HRP (Sigma) antibody for 90 min. Antibody was prepared in 1% BSA in TBS at a dilution of 1:5000. Following antibody incubation, cells were washed 3 times with 1% BSA (in TBS). Thereafter, cells were incubated with 200 µL of TMB-ELISA (Thermo Scientific) till the light blue color appeared and the signal was quenched by transferring 100 µL of colored solution to another 96-well plate containing 100 µL of 1 M $H_2SO_4$. Absorbance was measured at 450 nm using a multi-mode plate reader. For normalization of signal intensity, cell density was estimated by using a mitochondrial stain Janus green B. Afterward, TMB was removed and cells were washed 2 times with 200 µL of TBS. After washing, the cells were incubated for 15 min with 0.2% (w/v) Janus Green (Sigma). Excess stain was removed by washing with distilled water (3 times). The stain was eluted by adding 800 µL of 0.5 N HCl per well. 200 µL of eluted solution was transferred to a 96-well plate and absorbance was recorded at 595 nm. The signal intensity was normalized by calculating the ratio of A450/A595 values. For data normalization, ratio of A450/A595 value of pcDNA transfected cells reading was considered as 1 and calculated receptor expression with respect to pcDNA. Data were analyzed by using column in GraphPad Prism software.

**Confocal microscopy**

For visualizing βarr2 recruitment and its trafficking, HEK293 cells were transfected with 5 µg of $CXCR7^{WT}$ or $CXCR7^{E358A}$ along with 2 µg of βarr2-mYFP mammalian constructs using a polyethylenimine reagent (21 µL) in 10 cm plates. After 6 h, the transfected HEK293 cells in FBS-deficient DMEM media were now removed and replaced with 10% FBS-supplemented DMEM media (Gibco). Post 24 h, transfected cells were trypsinized and seeded onto poly-D-lysine (Sigma) precoated glass bottom confocal dishes (GenetiX) at a density of 1 million per plate. Simultaneously, with the same transfected cells another 24 well plate was seeded at 0.1 million per well for comparing the surface expression of $CXCR7^{WT}$ and $CXCR7^{E358A}$. After overnight culturing the cells in confocal dishes, cells were now starved in FBS-deficient

DMEM media for 4 h. Cells were visualized under the microscope and stimulated with CXCL12 (100 nM, Peprotech). Live cell microscopy was done on the cells and images were captured at multiple time points. The instrument used was Zeiss LSM 710 NLO confocal microscope where cells were housed on a temperature and $CO_2$-controlled platform with a motorized XY stage. A diode laser at 488 nm laser line was used for exciting mYFP and an emitted signal was detected with a 32×array GaAsP descanned detector (Zeiss). For all experiments, microscopic settings including laser intensity and pinhole slit were kept in the same range.

**Tango assay**

Tango assay measures βarr2 recruitment to the receptor. HTLA cell line, a derivative of HEK293 cell line stably expressing a tTA-dependent luciferase reporter gene and βarr2-TEV fusion gene was maintained in Dulbecco's Minimum Essential Media supplemented with 10% FBS, 2 μg $mL^{-1}$ puromycin and 100 μg $ml^{-1}$ hygromycin B at 37°C in 5% $CO_2$. Briefly, HTLA cells at a density of $3 \times 10^6$ cells per 10 cm plate were transfected with 7 μg of $CXCR7^{WT}$ or $CXCR7^{E358A}$ constructs. After 24 h of transfection, cells were trypsinized and seeded at a density of 0.05 million cells per well in a white 96-well polystyrene micro-plate. After another 16 h, cells were stimulated with varying doses of CXCL12 (0.32 μM to 0.1 pM). The ligand concentrations were prepared in incomplete media devoid of FBS. After incubation with ligand, media was aspirated and 100 μL of luciferin (0.5 mg $mL^{-1}$ in HBSS buffer) was added to each well and the plate was read for luminescence. The data was normalized with respect to the highest dose of ligand in $CXCR7^{WT}$ after basal correction, and then analyzed using nonlinear regression in GraphPad Prism software.

**HDX-MS analysis**

Wild-type βarr2 was prepared at a final concentration of 100 μM in a buffer consisting of 20 mM HEPES pH 7.4 and 150 mM NaCl. For peptide binding, 500 μM of C7pp2 was added to βarr2 and incubated at room temperature for 1 h. To initiate hydrogen/deuterium exchange, 2 μL of protein samples were mixed with 28 μL of $D_2O$ buffer (20 mM HEPES pH 7.4, 150 mM NaCl, and 10% glycerol in $D_2O$) and incubated on ice for 10, 100, 1000, or 10 000 seconds. At the indicated time points, the reaction was slowed down by adding 30 μL of ice-cold quench buffer (100 mM $NaH_2PO_4$ pH 2.01). Non-deuterated samples were prepared by mixing 2 μL of the protein sample with 28 μL of $H_2O$ buffer (20 mM HEPES pH 7.4 and 150 mM NaCl in $H_2O$) and subsequently quenching the reaction with 30 μL of ice-cold quench buffer. The quenched samples were subjected to online digestion by passing them through an immobilized pepsin column (2.1 mm × 30 mm) at a flow rate of 100 mL $min^{-1}$ using 0.05% formic acid in $H_2O$ as the mobile phase at 12°C. After the digestion process, the peptide fragments were collected on a C18 VanGuard trap column (1.7 mm × 30 mm) for desalting with 0.05% formic acid in $H_2O$. The proteins were subsequently separated using ultra-pressure liquid chromatography on an ACQUITY UPLC C18 column (1.7 μm, 1.0 mm × 100 mm) at a flow rate of 40 mL $min^{-1}$. The separation was achieved by employing an acetonitrile gradient generated by two pumps, initially starting with 8% B and gradually increasing to 85% B over a duration of 8.5 min. Mobile phase A consisted of 0.15% formic acid in $H_2O$ and mobile phase B comprised 0.15% formic acid in acetonitrile. To minimize the back exchange of deuterium to hydrogen, all components involved in the analysis, including the sample, solvents, trap, and UPLC column, were maintained at a pH of 2.5 and 0.5°C. Mass spectral analyses were performed with a Xevo G2 QTof equipped with a standard ESI source (Waters, Milford, MA, USA). The mass spectra were acquired in the positive ion mode in the range of m/z 100–2000 for 12 min. Peptides were identified in non-deuterated samples with ProteinLynx Global Server (PLGS) 2.4 (Waters, Milford, MA, USA). The following parameters were applied:

monoisotopic mass, non-specific for the enzyme while allowing up to 1 missed cleavage, MS/MS ion searches, automatic fragment mass tolerance, and automatic peptide mass tolerance. The searches were conducted with the variable methionine oxidation modification and the peptides were filtered with a peptide score of 6. To analyze the HDX-MS data, the deuterium content in each peptide was determined by measuring the centroid of the isotopic distribution using DynamX 2.0 (Waters, Milford, MA, USA). All measurements were performed in three independent experiments, and statistical significance was assessed using one-way ANOVA. Back-exchange levels were not corrected because the analyses compared different states. The details of HDX-MS data are described in Dataset S1.

### Supplementary figures

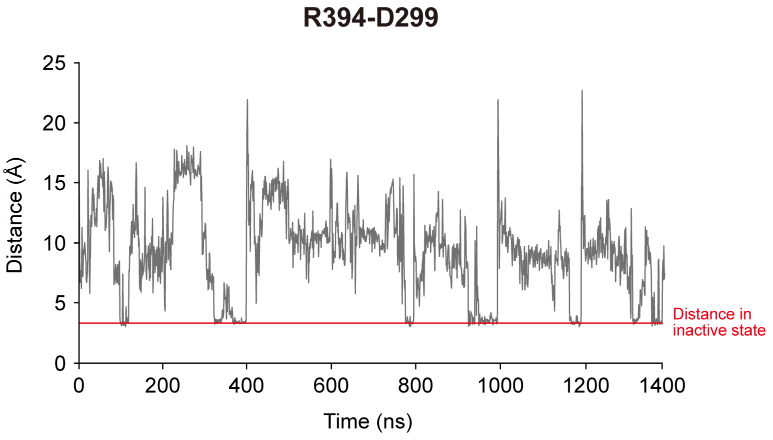


**Fig. S1.** Spontaneous inactivation of the C-tail observed during simulations of βarr2 with a manually displaced C-tail. The distance between R394 and D299 (forming polar interactions in the inactive state of βarr2) monitored within MD simulations (7 × 200 ns).

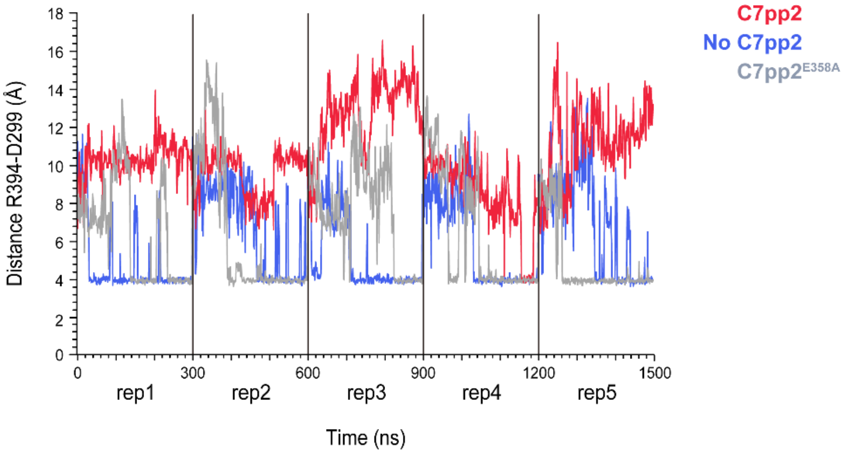


**Fig. S2.** The distance between R394 and D299 (forming polar interactions in the inactive βarr2 state) monitored within MD simulations (5 × 300 ns).

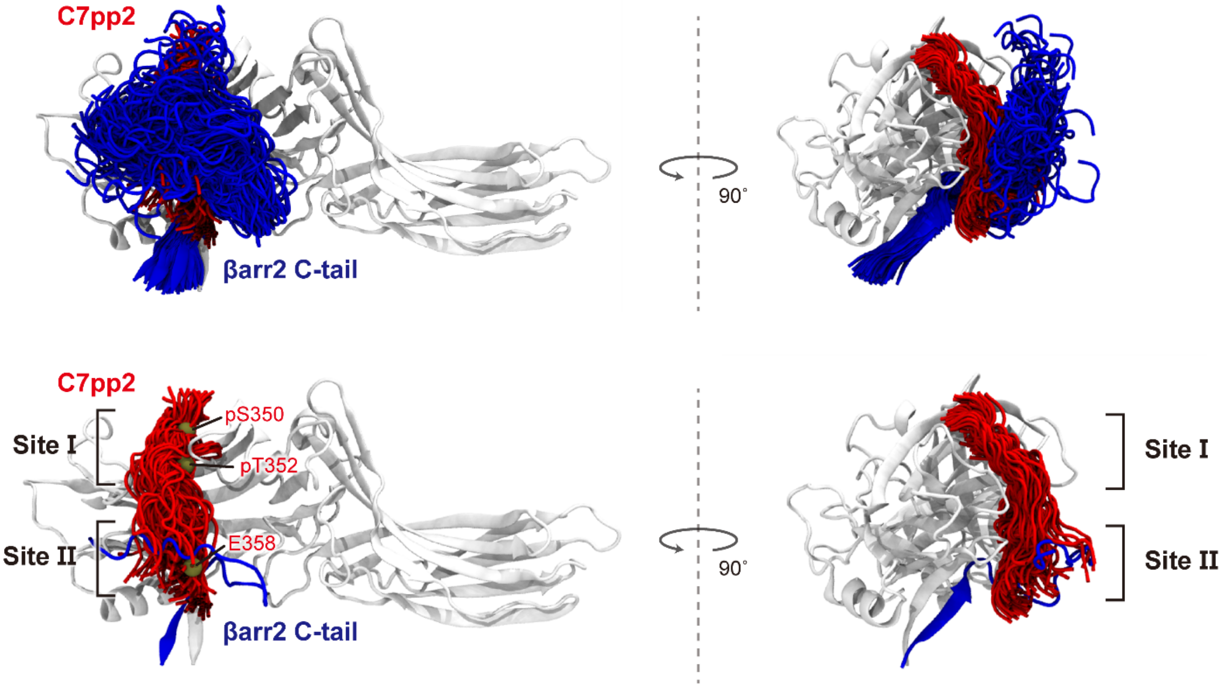


**Fig. S3.** Structural flexibility explored by the βarr2 C-tail (blue) and C7pp2 (red) explored in MD simulations (5 × 300 ns, 1 snapshot every 10 ns).

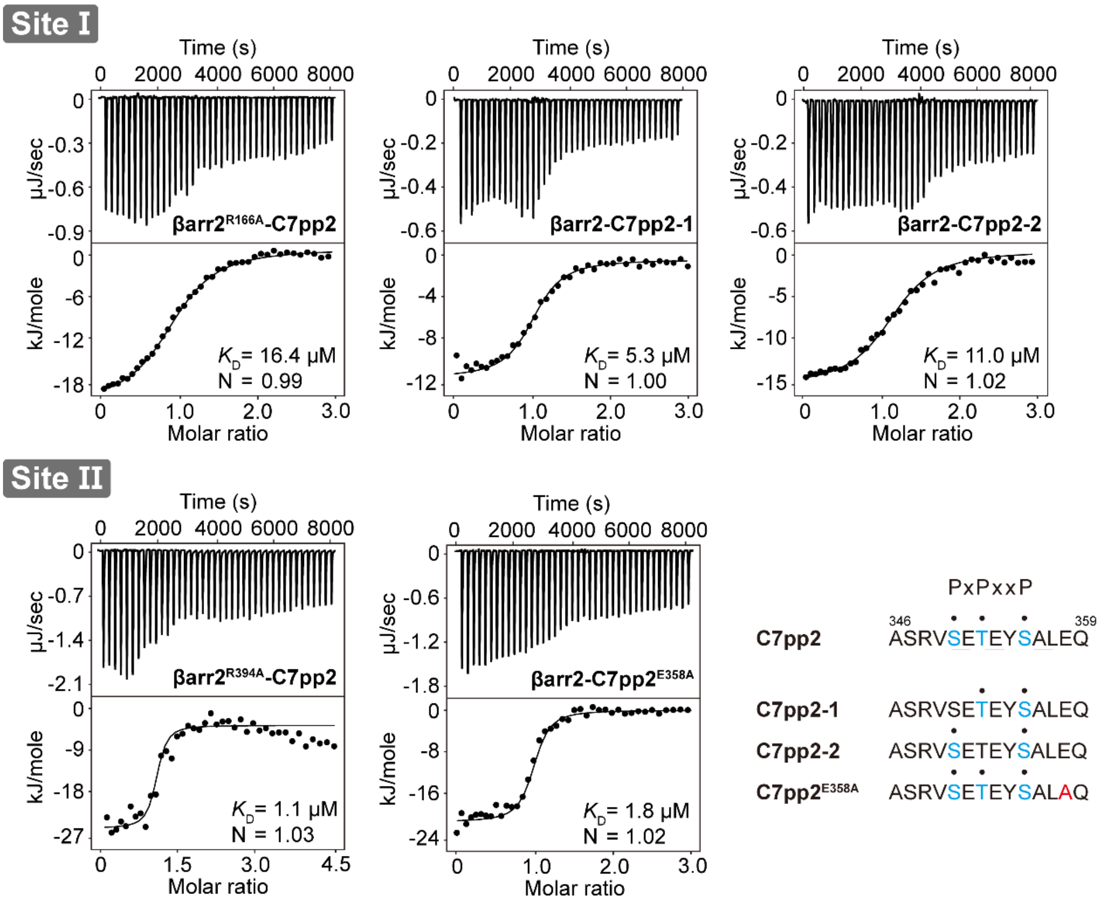


**Fig. S4.** The two binding interfaces of βarr2$_{IM}$-C7pp2 including site I and site II. ITC experiments showing the effect of mutations in the binding interfaces between βarr2 and C7pp2. The detailed sequences of C7pp2 mutants used for the ITC assay are shown in the lower right panel. Purified βarr2 was incubated with increasing peptide concentrations, and the binding parameters were calculated based on the dose-response curve.

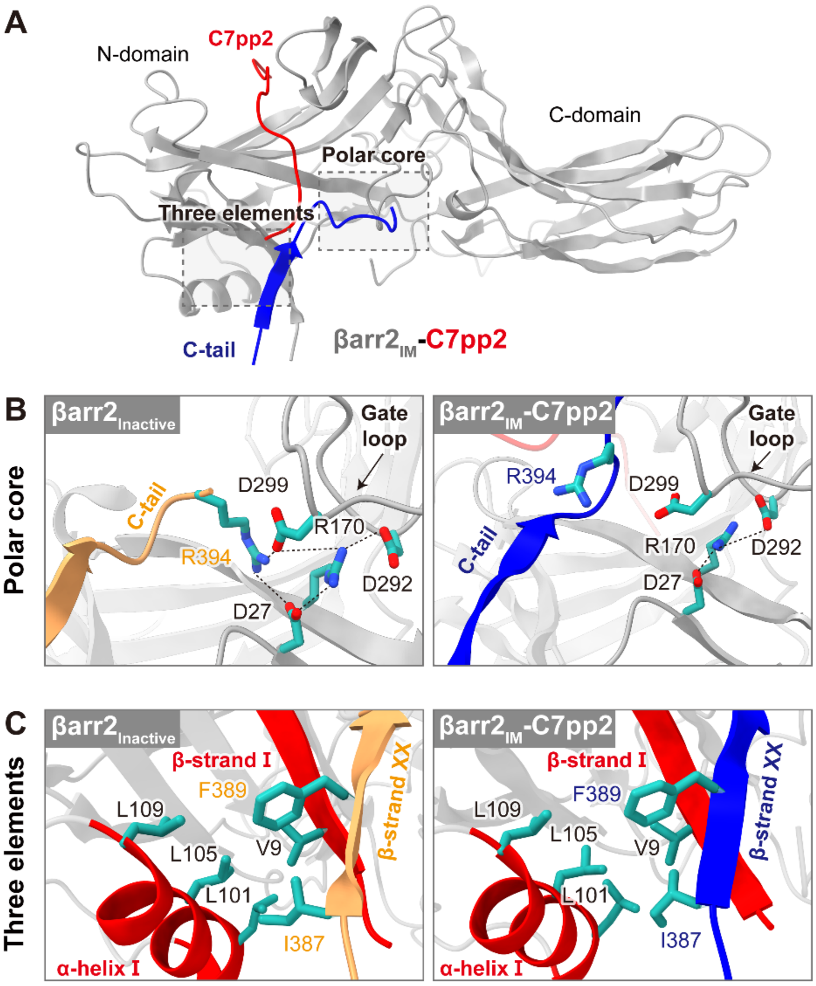


**Fig. S5.** Comparison of the inactive and intermediate conformations of βarr2. (A) Ribbon diagram of the C7pp2 bound to βarr2$_{IM}$. C7pp2 and the C-tail of βarr2 are shown in red and blue, respectively. The polar core and three elements are indicated in dotted squares. (B) Structural comparisons of the polar core in inactive βarr2 and βarr2$_{IM}$. Polar interactions are depicted as dotted lines. The polar core interactions are partially disrupted in the βarr2$_{IM}$-C7pp2 complex. (C) Structural comparisons of the three elements interactions in inactive βarr2 and βarr2$_{IM}$. The three hydrophobic elements stabilizing the basal state are highlighted in red (α-helix I and β-strand I), orange (β-strand XX in the inactive state), or blue (β-strand XX in the intermediate conformation).

#### Supplementary movie caption

**Movie S1 (separate file)**. Molecular dynamics simulation of the spontaneous inactivation of the βarr2 C-tail.

#### Supplementary dataset caption

**Dataset S1 (separate file).** The full profiles of HDX-MS data are provided in accordance with the recommendations of the international society of HDX-MS.
